# Growth couples temporal and spatial fluctuations of tissue properties during morphogenesis

**DOI:** 10.1101/2023.10.23.563640

**Authors:** Antoine Fruleux, Lilan Hong, Adrienne H. K. Roeder, Chun-Biu Li, Arezki Boudaoud

## Abstract

Living tissues display fluctuations – random spatial and temporal variations of tissue properties around their reference values – at multiple scales. It is believed that such fluctuations may enable tissues to sense their state or their size. Recent theoretical studies developed specific models of fluctuations in growing tissues and predicted that fluctuations of growth show long-range correlations. Here we elaborated upon these predictions and we tested them using experimental data. We first introduced a minimal model for the fluctuations of any quantity that has some level of temporal persistence or memory, such as concentration of a molecule, local growth rate, or mechanical property. We found that long-range correlations are generic, applying to any such quantity, and that growth couples temporal and spatial fluctuations, through a mechanism that we call ‘fluctuation stretching’ — growth enlarges the lengthscale of variation of this quantity. We then analysed growth data from sepals of the model plant Arabidopsis and we quantified spatial and temporal fluctuations of cell growth using the previously developed Cellular Fourier Transform. Growth appears to have long-range correlations. We compared different genotypes and growth conditions: mutants with lower or higher response to mechanical stress have lower temporal correlations and longer-range spatial correlations than wild-type plants. Finally, we used theoretical predictions to merge experimental data from all conditions and developmental stages into an unifying curve, validating the notion that temporal and spatial fluctuations are coupled by growth. Altogether, our work reveals kinematic constraints on spatiotemporal fluctuations that have an impact on the robustness of morphogenesis.

**Significance Statement:** How do organs and organisms grow and achieve robust shapes in the face of subcellular and cellular variability? In order to address this outstanding mystery, we investigated the variability of growth at multiple scales and we analysed experimental data from growing plant tissues. Our results support the prediction that tissue expansion couples temporal memory of growth with spatial variability of growth. Our work reveals a constraint on the spatial and temporal variability of growth that may impact the robustness of morphogenesis.

## INTRODUCTION

The impact of noisy perturbations on organism development is the subject of active research [1]. Fluctuations – the random spatial and temporal variations of tissue properties around their reference values – have been observed at multiple scales, from cytoskeleton [2] to cell [3] and tissue [4]. In the fruit fly, for example, actomyosin pulses were shown to cause fluctuations of cell shape [5–7], while fluctuations of the position of cell junctions were found to favor cell rearrangements during tissue extension [8, 9]. It was proposed that fluctuations are required for symmetry breaking and pattern formation during development [10, 11] or for cells and tissues to sense their neighbourhood [12]. Fluctuations in gene expression or morphogens seems particularly important for cell differentiation. Fluctuations in gene transcription seem required for the maintenance of pluripotency [13, 14], and specific properties of fluctuations are a signature of cell differentiation [15–18]. Nevertheless, the robustness of tissue patterning appears sensitive to fluctuations in molecule concentrations [19, 20]. Fluctuations in growth induce mechanical stress [12, 21–23] because, for instance, cells with higher growth rate exert forces on neighbouring cells, which may sense and respond to such mechanical stress. Robust development of the fruit fly wing partially relies on cell competition, i.e. on mismatch of growth rates between cells, and on the ensuing modulation of proliferation and apoptosis [24, 25]. In this context, it is important to understand whether fluctuations of a cell affect its local neighbourhood or the whole tissue. Here, we analysed the spatial structure of fluctuations in experimental data from growing tissues.

Recent models of tissue mechanics and growth accounted for temporal and spatial fluctuations of growth and investigated their role in robustness of morphogenesis [26–28]. Temporal fluctuations are characterised by their degree of persistence, quantified with the persistence time (or correlation time), the characteristic time over which memory of previous fluctuations is lost. It could be the time needed for remodelling of the cytoskeleton or of the extra-cellular matrix (in animals) / the cell wall (in plants). Spatial fluctuations are characterised by their degree of spatial consistency, quantified by the correlation length, the characteristic length over which cells (or subcellular domains) behave similarly, or by cell-to-cell variability over a small neighbourhood. For instance, the shape of a plant organ was found to be less robust in a mutant with lower cell-to-cell variability [26]. However, spatial fluctuations may have a more complex structure. Indeed, theoretical models of the expanding universe [29, 30] and of growing tissues [27, 28] predicted long-range spatial correlations, i.e. a significant level of correlations between fluctuations of two distant parts of the system; accordingly, growing systems are expected to exhibit fluctuations at multiple scales. Here we focus on the underlying mechanism, which we call fluctuation stretching – the increase in the lengthscale of fluctuations of a tissue property or of the concentration of a molecule, due to tissue expansion.

To assess the experimental relevance of this mechanism, we analyzed growth fluctuations in the model plant *Arabidopsis thaliana*. We considered the sepal, the green leaf-like organ that protects a flower prior to its opening. We characterised sepals from wild-type individuals in different culture conditions as well as mutant plants. We considered *spiral2* and *katanin* mutant plants since they were found to be less robust to variability in the number of trichomes (epidermal hair-like cells) than wild type plants [31], suggesting a greater impact of cellular scales on organ ones. The lack of SPIRAL2 and KATANIN function led respectively to stronger [31–33] and weaker [31, 32, 34] cortical microtubule co-alignment and reorientation in response to mechanical stress [35, 36]. Microtubules guide the deposition of cellulose fibers in the cell wall (the plant extra-cellular matrix) [37]. Cellulose fibers being the main load-bearing component of the cell wall, the response of microtubules to mechanical stress is generally considered as a mechanical feedback on growth and *spiral2* and *katanin* as mutants with altered feedback.

In this Article, we first present a simple model for fluctuation stretching. We estimate spatial and temporal correlations of tissue growth fluctuations in Arabidopsis sepals using previous live imaging data [31, 32] and the Cellular Fourier Transform (CFT) [38]. We investigate how correlations vary within and between datasets and we test the relevance of fluctuation stretching.

## RESULTS

### A minimal models predicts the stretching of fluctuations in growing tissues

Fluctuation stretching, the enlargement of the length-scales of fluctuations by medium expansion, was predicted by different models of expanding media, the early universe [29, 30] and living tissues [27, 28]. Here we introduce a minimal model for fluctuation stretching. For a primarily mostly interested in experimental data, Eq. 2 is the main theoretical result that we test in growing sepals.

We consider a variable property Φ that is defined on a tissue growing isotropically at average rate 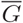 and that depends on position vector *x* and time *t*. This variable Φ could reflect gene expression, signalling, metabolism, cell size, or cell growth, for instance. We assume that (i) Φ is inherited through tissue growth, so that it is advected (transported) by the average growth velocity 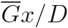 (*D* is the space dimension: *D*=1 in Figures 1-2 and *D* = 2 for a thin organ like the sepal), (ii) Φ relaxes to its average value ⟨Φ⟩ with a characteristic memory (persistence/correlation) time *τ*, and (iii) Φ is subject to a source of noise *ξ*(*x, t*) that is random in space and time. As a consequence,

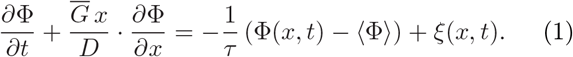

In this equation, the first term is the temporal derivative of Φ(*x, t*). The second term (in right-hand side) represents the effect of tissue expansion, i.e. advection by growth, and contains the spatial derivative of Φ (the dot · stands for the vectorial product, which reduces to a multiplication for *D* = 1). The third term (left-hand side) describes relaxation (loss of memory) of Φ.

**FIG. 1.**
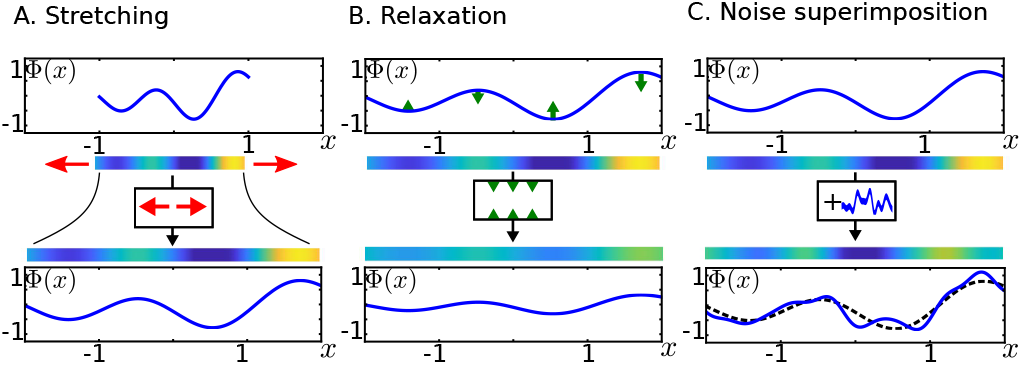
Distinct effects of tissue expansion, time relaxation (loss of memory), and noise source on the spatial pattern of a tissue property. The figure shows initial spatial patterns and their temporal evolution under the three mechanisms. The variable property Φ(*x*) is plotted as a function of position *x* and shown in colorscale (blue and yellow for low and high values, respectively) along a strip standing for the growing tissue. **A** Tissue expansion induces fluctuation stretching, defined as the enlargement of the length-scales of fluctuations. **B** Relaxation associated with loss of memory induces a decay in the amplitude of fluctuations (depicted by green arrows). **C** Noise causes the superimposition of new fluctuations on the preceding pattern (represented by a dashed line in the lower panel). We schematically represent stretching, relaxation, and noise superimposition by function block diagrams containing horizontal red arrows, vertical green arrows, and a noisy signal, respectively. These block diagrams are used in Fig. 2.

The consequences of tissue expansion, loss of memory (time persistence), and noise on the variations of Φ are schematized in Fig. 1, for one time step. Tissue expansion induce ‘fluctuation stretching’, i.e. enlarges the length-scales of spatial variations (panel **A**). Time persistence determines how fast fluctuations relax toward their reference level (**B**). Noise superimpose new fluctuations on the preceding pattern (**C**).

When iterated over time, fluctuation stretching and noise give rise to multiscale fluctuations, while the degree of time persistence (or memory level) controls how far fluctuations extend toward large space-scales. This is illustrated in Fig. 2**A**. in three regimes: for full, intermediate, and vanishing time persistence. For full time persistence 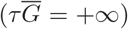 the pattern is stretched, increasing its the lengthscale of variations of Φ and fluctuations are added at small scale. For intermediate time persistence 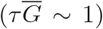, the same process occurs but the preexisting pattern is attenuated due to relaxation. In the absence of temporal persistence (*τ* = 0), the preceding pattern disappears and only the newly superimposed noise remains. Mathematically, the solutions to Eq. 1 take the form 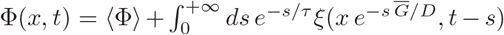 (see Supplementary note, for details). The integral indicates the superimposition while the exponential factor *e*^−*s/τ*^ accounts for time relaxation or loss of memory. Fluctuation stretching corresponds to the exponential factor 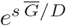 applied to the spatial variation of the noise.

**FIG. 2.**
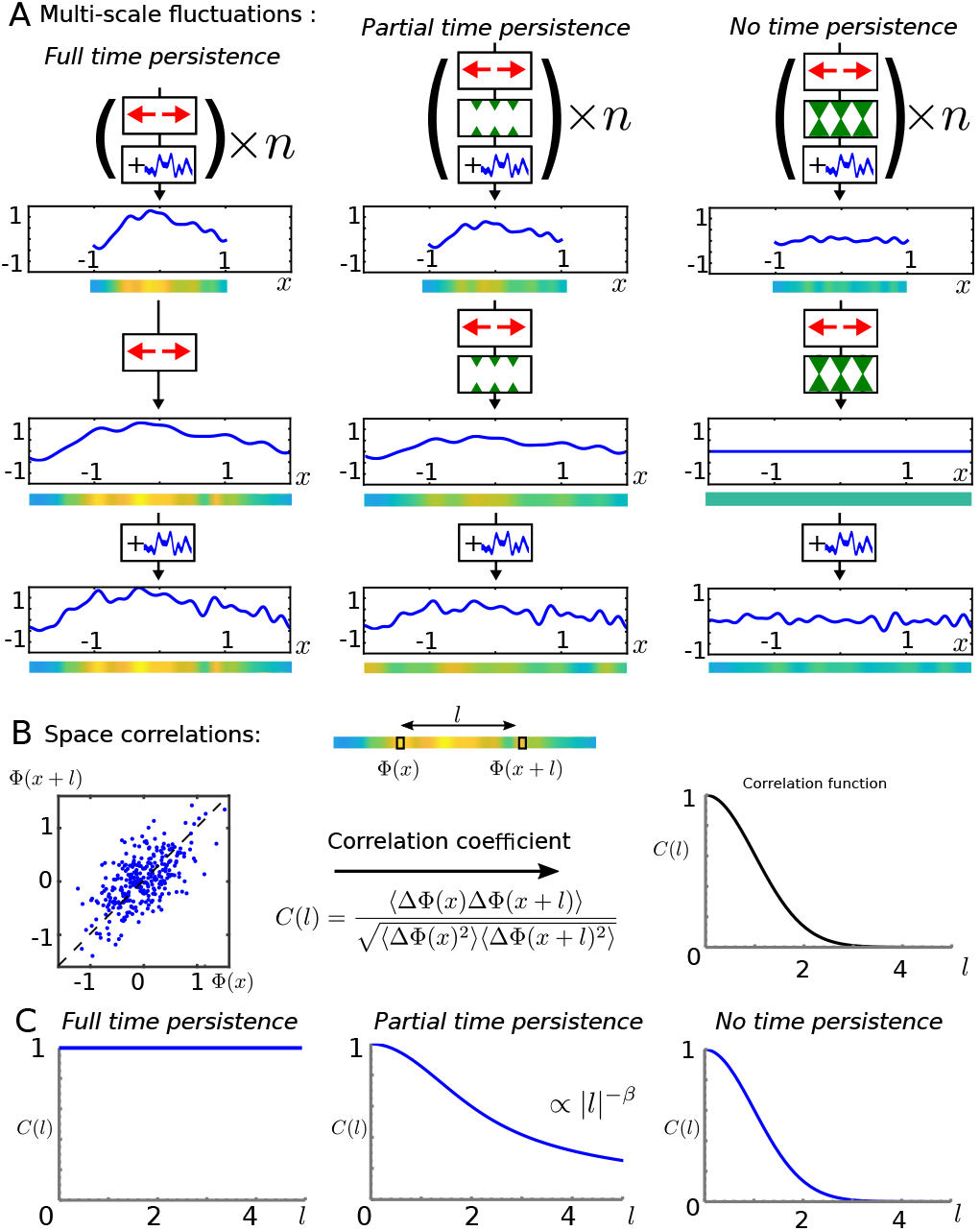
Multiscale fluctuations as a consequence of fluctuation stretching. Spatial correlations of tissue properties depend on the level of temporal persistence of fluctuations. Three levels of temporal persistence are considered: full (no time relaxation), intermediate (moderate relaxation), and none (instantaneous relaxation). **A** Spatial pattern resulting from the iteration of fluctuation stretching, relaxation, and noise, schematically represented by function block diagrams in series, as defined in Fig. 1; patterns are represented under the form of plots and color stripes as in Fig. 1. Top: patterns after *n* iterations; middle: patterns after one additional iteration of stretching and (if appropriate) relaxation; bottom: patterns after one additional superimposition of noise. **B** Quantification of spatial correlations. Top: This involves comparing the values of the variable at positions *x* and *x* + *l*, as illustrated in the colored strip. Left: Typical scatter plot showing Φ(*x* + *l*) as a function of Φ(*x*) for multiple values of *x*. Middle: *C*(*l*) is defined as the correlation coefficient between Φ(*x*+*l*) and Φ(*x*); ⟨ ⟩stands for the statistical average of the expression between brackets and ΔΦ(*x*) = Φ(*x*) − ⟨ (Φ(*x*) ⟩. Right: the correlation *C*(*l*) as a function of the distance *l*. **C** Spatial correlation function *C*(*l*) for full, partial, and no time persistent fluctuations. Models predict that the space correlation function is a power-law of *l, C*(*l*) ∝ *l*^−*β*^.

The space correlation function, *C*(*l*), is the pairwise correlation between the values Φ(*x*) and Φ(*x* + *l*) of the variable Φ at positions distant of length *l*, as illustrated in Fig. 2 **B**. *C*(*l*) generally decrease with the distance *l*: for *l* = 0, Φ(*x*) = Φ(*x* + *l*) and so the correlation is complete, *C*(0) = 1, while at large distance *l*, Φ(*x* + *l*) is expected to be independent of Φ(*x*) and the correlation vanishes as illustrated in the plot on the right of panel **B**. In our minimal model, the correlation function takes the form 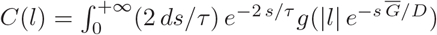, assuming the permanent noise source *ξ*(*x, t*) = 0 has zero mean and correlation function ⟨ *ξ*(*x, t*)*ξ*(*x* + *l, t* + *s*) ⟩ proportional to *δ*(*s*)*g*(*l*) (*δ* is the Dirac distribution, see Supplementary note, for details). Here again *C*(*l*) appears as a weighted sum of the space correlation function *g* of the noise source stretched at different spatial scales. The correlation function *g* is assumed to have a correlation length *𝓁* that sets the reference scale for spatial variations of Φ; *𝓁* cannot be assumed to be zero without causing issuess of mathematical convergence. In practice, we took 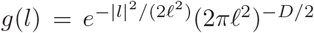. Because of fluctuation stretching, space correlations functions for time persistent fluctuations are predicted to be long-ranged *i*.*e*. to have their tails which follow a power law ∝ *l* ^−*β*^. As shown in the Supplementary note, this can be made explicit by rewriting the space correlation function 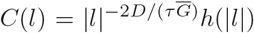 where the increasing function 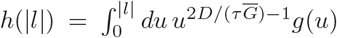, reaches an asymptotic value when |*l*| becomes large compared to the correlation length *𝓁* of *ξ*. Therefore, the correlation function *C*(*l*) of the variable of interest Φ mostly behaves as a power-law *C*(*l*) *∼ l*^−*β*^ of exponent

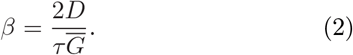

This scaling law indicates that the values of the variable Φ considered in two distant points decorrelate slowly as their distance is increased, which reflects the fact that fluctuations are a superimposition of patterns with different spatial lengthscales. *β* estimates this spatial decrease in correlations, the higher the memory (the larger 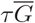), the higher correlations between distant regions. Fig. 2 **C** show the space correlation functions for full, partial, and no memory. Full temporal persistence is simply the limit where the persistence time is infinite, leading to an accumulation of fluctuations at large lengthscales. The weight of large scaled fluctuations continuously increases so that the correlation function tends toward a constant. In contrast, in the absence of temporal persistence, spatial correlations vanish beyond the correlation length of the noise. Hereafter, we tested this prediction using previous experimental data about growing plant organs.

### Live imaging and spectral analysis provide estimates for spatiotemporal correlations of cell growth

Next we aimed at a quantitative description of spatial and temporal correlations of growth fluctuations in expanding tissues. We used experimental data where sepals were imaged live to track morphogenesis over time, with similar culture and imaging conditions [31, 32]. We examined whether fluctuations stretching applies to cell areal growth rate. Each sepal was imaged at multiple times, labeled *t* = 0, 1, 2, … and separated by 24 hours intervals as illustrated by Fig. 3**A**, which shows an example of cells segmented in a sepal, at three successive time steps *t, t* + 1 and *t* + 2. Growth was defined from cell surface area at successive time steps. Fig. 3**B** shows cell areal relative growth rate *G*_*i,t*_ and *G*_*i,t*+1_ from *t* to *t* + 1 and from *t* + 1 to *t* + 2 respectively, deduced from segmentation of sepals into cells, as showed in panel **A** and mapped on the reference tissues at *t* and *t* + 1, respectively. When a cell has divided between *t* to *t* + 1, we used the total surface area of its daughter cells at *t* + 1 to define *G*_*i,t*_, see Datasets ans Methods for details.

**FIG. 3.**
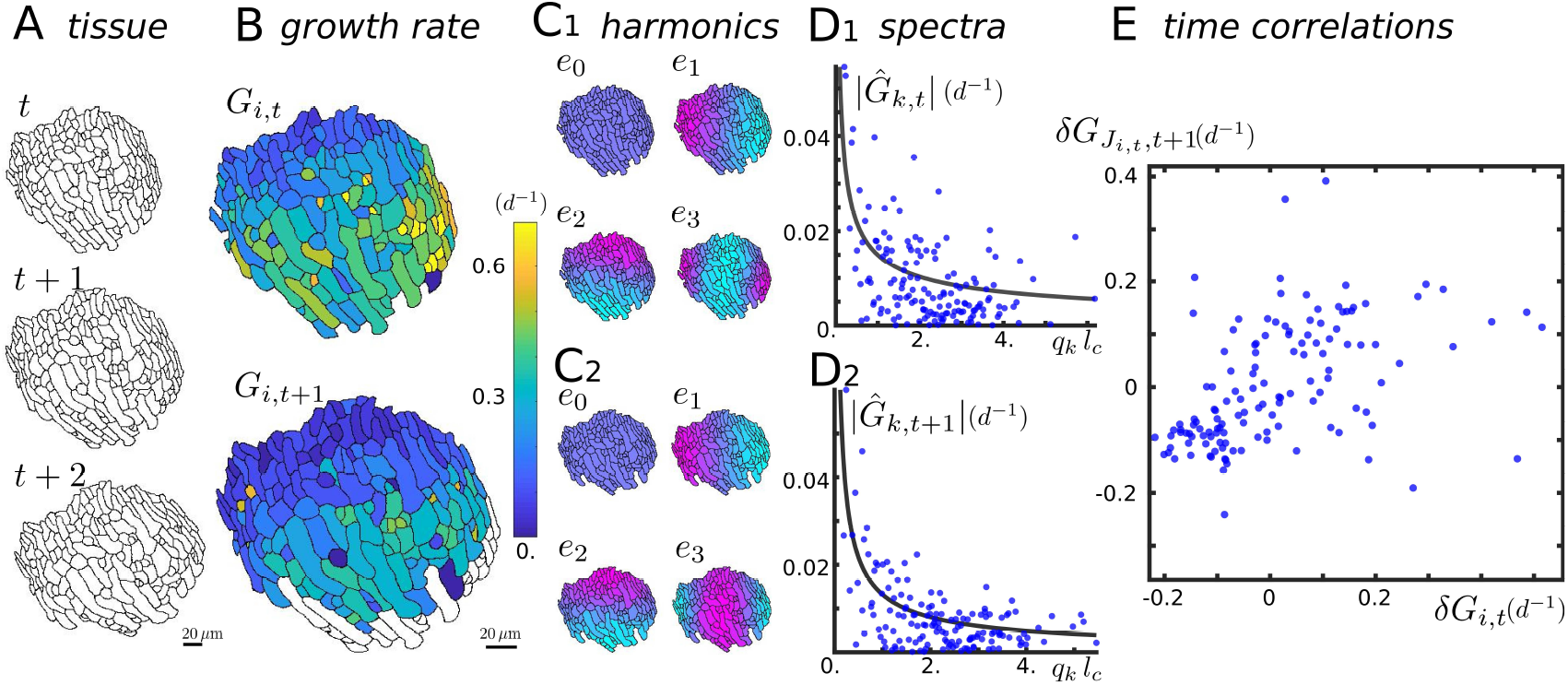
Quantification of spatial and temporal fluctuations in cell growth. Day (*d*) is used as a unit of time. **A** Three snapshots of a plant tissue (abaxial sepal epidermis from wild-type plant) taken at one-day intervals. Black lines represent cell contours. **B** Heatmaps of relative areal growth rate between times *t* and *t* + 1, *G*_*i,t*_, and between *t* + 1 and *t* + 2, *G*_*i,t*+1_ for cell #*i*. A growth rate of 1*d*^−1^ corresponds to a relative increase of area of 100% in 1 day. Growth rate of white cells could not be computed because they were not imaged at *t* + 2. **C**_1_**-C**_2_ The first 4 harmonics *e*_*k*_ (*k* = 0, 1, 2, and 3) of the Cellular Fourier Transform (CFT) of the tissue at *t* and *t* + 1 (the white cells in **B** are not included), represented by a cyan (low value) to magenta (high values) color scheme. The harmonics *e*_*k*_ generalise sinusoidal waves and can be used to decompose the growth fields *G*_*i,t*_ and *G*_*i,t*+1_ into their respective CFTs 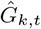 and 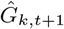. **D**_1_**-D**_2_ Fourier spectra (blue dots) correspond to the absolute values 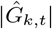 and 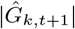 of the CFTs and are shown as function of the wavenumber *q*_*k*_ of the harmonics *e*_*k*_. Wavenumbers were non-dimensionalised using mean cell size *l*_*c*_. A representative power-law (solid line) 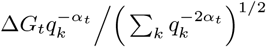 was obtained as explained in the text. Each spectrum is then characterised by two numbers, the standard deviation of cell growth Δ*G*_*t*_ and the spatial exponent of spatial correlations, *α*_*t*_. Here *α*_*t*_ = 0.54 ± 0.08 (± standard error of the mean), *α*_*t*+1_ = 0.71 ± 0.08, Δ*G*_*t*_ = 0.157 ± 0.012 *d*^−1^ and Δ*G*_*t*+1_ = 0.134 ± 0.012 *d*^−1^. **E** For temporal analyses, detrended areal growth rate *δG*_*i,t*_ was computed as the excess areal growth rate of a cell with respect to a local neighborhood. The coordinates of each blue dot are the detrended growth *δG*_*i,t*_ of a cell *i* between *t* to *t* + 1 (horizontal axis) and the detrended growth 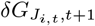 of the set *J*_*i,t*_ of its daughters between *t* + 1 and *t* + 2 (horizontal axis). The degree of growth temporal correlation is quantified by the value of the Kendall correlation coefficient, here Γ_*t*_ = 0.400 ± 0.052 (± standard error). Two outliers were excluded from the plot to improve the readability of the figure.

To dissect spatial variations of growth in the tissue, we used the Cellular Fourier Transform (CFT) [38]. The CFT consists of decomposing the signal into a linear combination of ad hoc harmonics that account for the subdivision of the tissue into cells of variable size and shape. These harmonics are the equivalent of sinusoidal waves in an infinite continuous medium. The *k*-th harmonic, *e*_*k*_, has wavenumber *q*_*k*_, and varies on a lengthscale that decreases with the rank *k*. The CFT coefficients 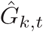 give the weights with which cell relative areal growth is decomposed into the harmonics *e*_*k*_. The Fourier spectrum is obtained by plotting the amplitude 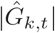 as a function the corresponding wave number *q*_*k*_. This spectrum is well-suited to describe fluctuations of *G* at multiple scales.

We investigated spatial correlations from Fourier spectra such as those shown in Fig. 3.**D**. The amplitudes of spectra appear significantly higher for low wave numbers, suggesting long-range correlations. To further test this, we sought a characteristic lengthscale for fluctuations and we considered the smallest index *K* for which 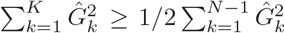, so as to quantify the repartition of fluctuations between low and large scales. If fluctuations were short-ranged, then the ratio of largest to characteristic wavenumbers, *q*_1_*/q*_*K*_, would be a good estimate of the ratio of correlation length to sample size, and would therefore be small compared to 1. In contrast, we found the ratio *q*_1_*/q*_*K*_ to be 0.54 on average (standard deviation 0.29 and range 0.086 – 1, over all study samples), indicating long-range correlations. This qualitative agreement with the predictions of the minimal model prompted us to use power-laws to represent Fourier spectra. We note that the prediction *C*(*l*) *∼ l*^−*β*^ corresponds to a spectrum scaling like *q*^−*α*^, with *α* = 1 − *β/*2 (see section Datasets and Methods). Although the limited range of wavenumbers did not allow us to test the power-law behavior, we obtained a representative power-law as follows. As the CFTs can be positive or negative, we assumed each CFT to follow a Gaussian distribution of zero mean and of standard deviation *σ*_*k,t*_, which was fitted to the equation 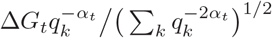. Each spectrum is then characterised by two numbers, its amplitude Δ*G*_*t*_ and its exponent, *α*_*t*_. The specific choice made for the fit is such that, following the Parseval theorem, Δ*G*_*t*_ measures the standard deviation of growth while *α*_*t*_ measures its spatial correlations. We used statistical inference to estimate *α*_*t*_ and Δ*G*_*t*_. The scaling exponent, *α*_*t*_, is expected to vary between 0 and 1, which correspond to short-range and to extremely long-range correlations, respectively. We found *α*_*t*_ to approximately range between 0.1 to 0.9, indicating large differences between samples and time points in terms of range of correlations (but see below for the comparison between genotypes). We found the standard deviation of growth Δ*G*_*t*_ to range between 0.1 and 0.6 *d*^−1^, values that are of the order of half the growth rate of a sample averaged over all cells between two time points, indicating relatively strong fluctuations of cell growth rate.

The temporal resolution (1*d*) and the number of consecutive images of a sample (3 to 7) were in general too low to compute persistence time from experimental data. We therefore estimated temporal persistence of growth using correlation coefficients. We considered the correlations between relative areal cell growth *G*_*i,t*_ from *t* to *t* + 1 and 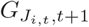 from *t* + 1 to *t* + 2, where the set *J*_*i,t*_ in subscript contains the labels of all daughters of cell *i* at time *t* and 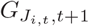 is their areal growth rate, see section Datasets and Methods for details. To avoid any bias due to overall gradients in growth rate [32], we computed detrended cell growth *δG*_*i,t*_ by substracting from the areal growth rate of a cell the average areal growth in a local neighborhood, see Supplementary note. The scatter plot in Fig. 3**E** of 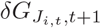 as a function of *δG*_*i,t*_ shows that growth is relatively persistent in time: For instance cells that grow more than their neighbors between *t* and *t* + 1 tend to remain so between *t* + 1 and *t*+2. We quantified temporal correlations of growth using Kendall’s correlation coefficient, Γ_*t*_, because it is based on the rank of data and is less sensitive to outliers than the more classical rank-based Spearman correlation coefficient [39]. Over all sepals and time points considered, Γ_*t*_ approximately ranges from 0.1 to 0.6. Almost all values of Γ_*t*_ were positive, while the negative values of Γ_*t*_ were not significantly different from zero (see below), indicating that, in general, growth is persistent over a time comparable to experimental time resolution (1*d*).

We thus obtained a minimal set of parameters to describe growth fields and their fluctuations: average growth rate,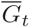, extent (exponent) of spatial correlations, *α*_*t*_, amplitude of spatial correlations, Δ*G*_*t*_, and temporal correlation coefficient Γ_*t*_. Next, we analysed differences and common features between sepals based on this minimal set of parameters.

### Temporal and spatial correlations of cell growth vary across genotypes and culture conditions

We analyzed growth fluctuations in several genotypes and culture conditions. As explained in the introduction, we chose to focus on mutants affected in responses to mechanical stress, *spiral2* (*spr2*) and *katanin* (two alleles, *bot1* and *mad5*), in addition to wild-type plants. We analyzed sepals from 4 genotypes in 2 culture conditions and at different developmental stages. In order to enable the comparison between several sepals that were imaged starting from different stages, we temporally aligned live imaging sequences along a common time frame using sepal width, building upon the approach developed in [40], see Datasets and Methods. The parameters that characterise growth fields in all these sequences are shown in Fig. 4.

**FIG. 4.**
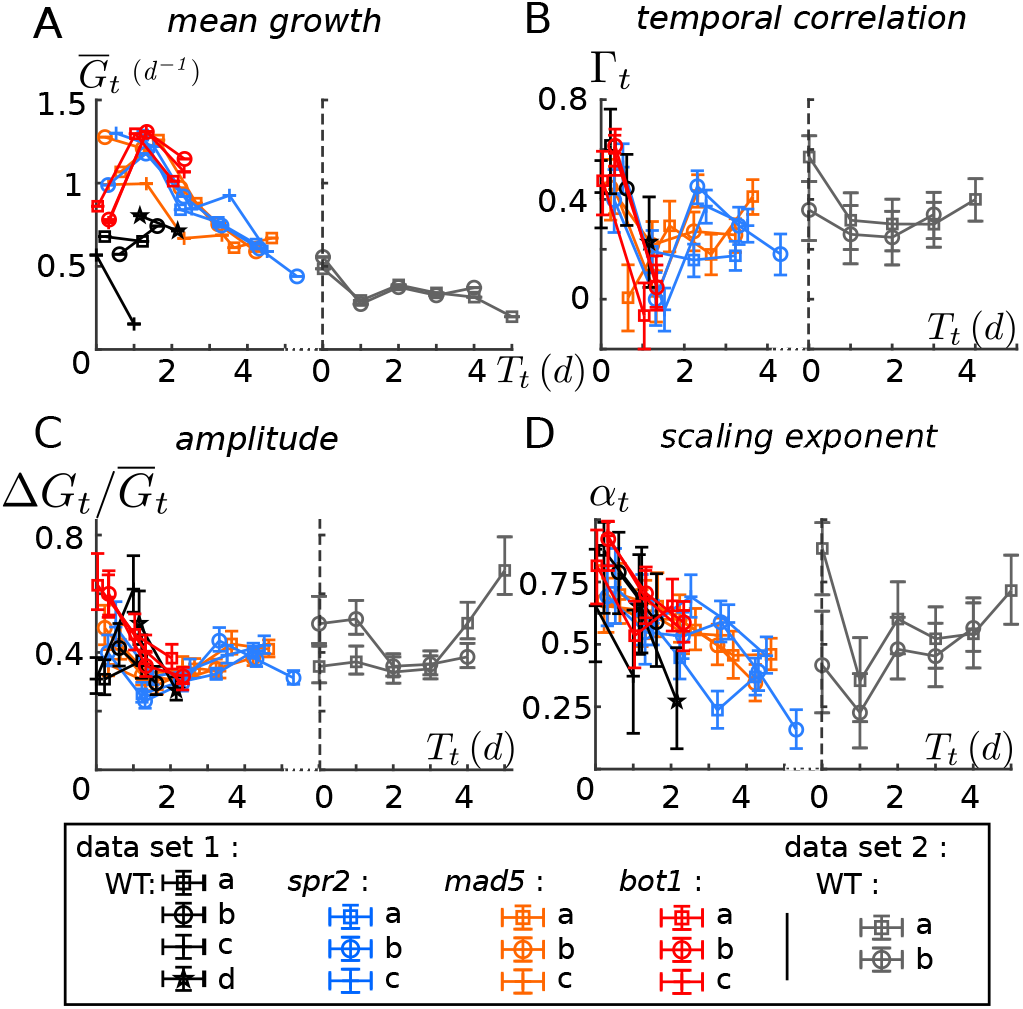
Parameters that characterise growth fields in sepals from wild-type and mutant plants. The sequences were temporally aligned and parameters are shown as a function of the synchronized time *T*_*t*_. **A** Growth rate averaged over the tissue 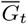. **B** Temporal correlation coefficient Γ_*t*_. **C** Dimensionless amplitude of the Cellular Fourier Transform (CFT) 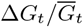 (also coefficient of variation of growth). **D** Scaling exponent of the CFT *α*_*t*_. The two datasets correspond to two slightly different culture conditions. Black, blue, orange and, red symbols/lines correspond respectively to wild-type, *spr2* mutant, *mad5* mutant, and *bot1* mutant from the first dataset, while gray symbols/lines correspond to wild-type plants from the second set. Error bars indicate the 90% confidence intervals; error bars are not shown in A because they are comparable to symbol size.

We first noticed a significant variability within and between genotypes/conditions and trajectories that seem heterogeneous in time. Some of this variability might be due to experimental constraints; for instance, the imaged regions of sepals varied in time and between individuals. We nevertheless observed a few trends that hold for several genotypes and conditions. Mean growth rate (panel **A**) decreases in time for trajectories that are long enough (*spr2, mad5* and wild-type in dataset 2), which is a general trend in organ morphogenesis. Temporal correlations (panel **B**) decrease between the first and the second time point, possibly associated with the strong decrease in growth anisotropy observed after the second time interval [32]. The relative amplitude of growth fluctuations (panel **C**) decreases for the first stages in mutants before stabilizing around 0.4. The extent of spatial correlations (panel **D**) tends to decrease with time in dataset 1.

In order to quantify differences induced by mutations or culture conditions, we used wild-type plants from dataset 1 as a reference and we estimated the shift in growth parameters between the reference and other genotypes or culture condition, see Fig. 5. As the amount of information available varied with genotype, culture condition, or temporal stage, we developed a method that enables a consistent comparison of differences by taking into account developmental stages, see Datasets and Methods for details. Briefly, we considered all pairs formed by a reference sepal (wild-type from dataset 1) and another sepal. We computed the shift between a reference sepal to another sepal at a given temporal stage and we averaged shifts over time and sepal pairs to obtain a mean shift, shown in Fig. 5 for all comparisons. This mean shift can be understood as the representative vertical difference between reference wild-type curves and mutant or dataset 2 curves from Fig. 4. We then estimated the standard error of these shifts, which results from the uncertainties of both reference sepals (wild-type from dataset 1) and sepals of the condition of interest.

**FIG. 5.**
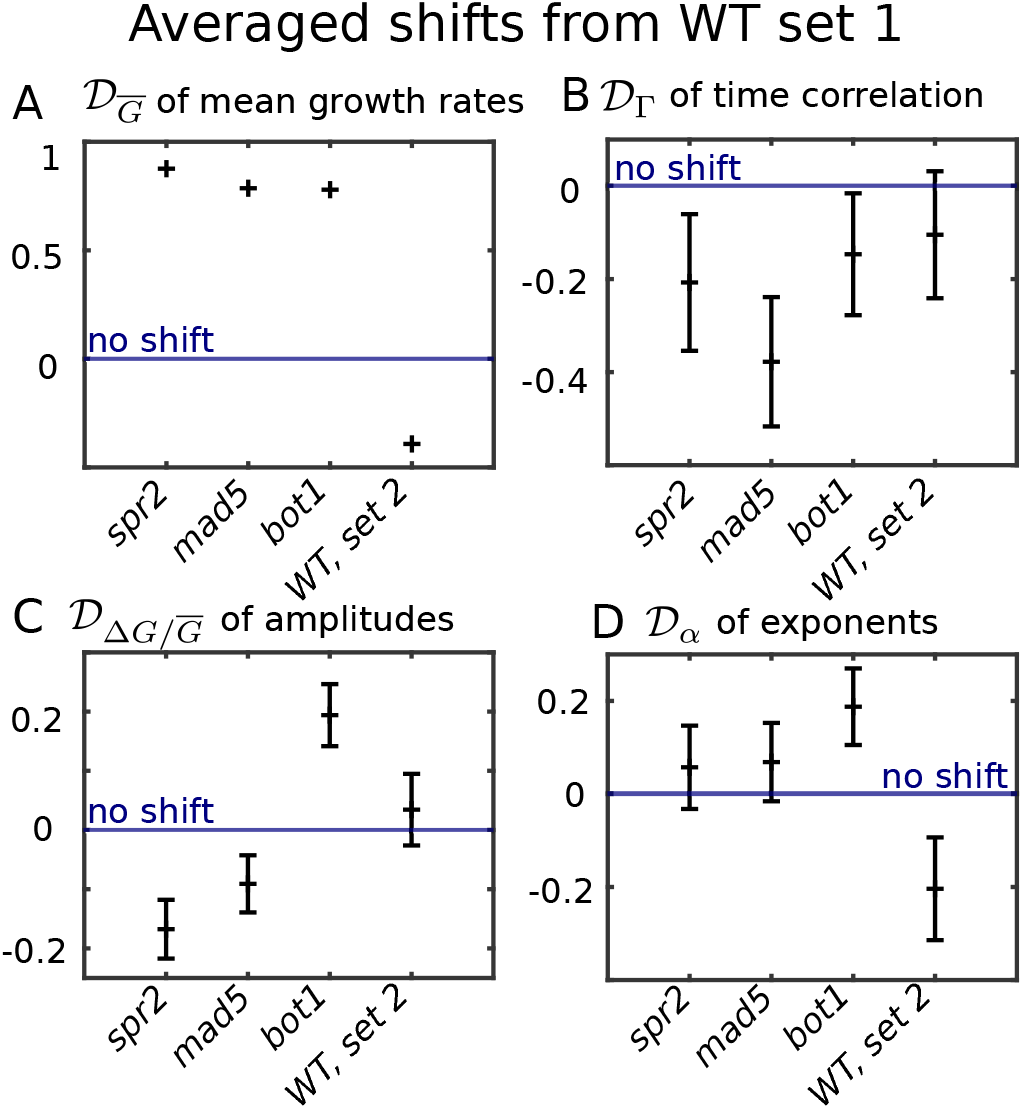
Differences in growth parameters due to mutations or to change in culture conditions. Data are shown for mutants from dataset 1 and wild-type (WT) from dataset 2; wild-type from dataset 1 was used as a reference in all cases. Symbols show the mean shifts 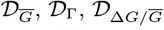 and 𝒟_*α*_ of : **A**, growth rates averaged over sepals, 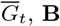, **B**, temporal correlation coefficients, Γ_*t*_, **C**, dimensionless amplitudes of growth fluctuations, 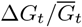, and **D**, exponents quantifying spatial extents of growth fluctuations, *α*_*t*_, respectively. Symbols and errors bars correspond to the mean and standard error of the difference, respectively; error bars correspond to the errors on the shifts 𝒟_Φ_ computed from the error on the data of interest (mutants or WT dataset 2) and on the reference one (WT dataset 1).

In wild-type, datasets 1 and 2 do not differ in temporal correlations (panel **B**) and amplitude of fluctuations (Fig. 5.**C**) within the range of uncertainty on these parameters. Average growth rate (Fig. 5.**A**) and extent of spatial correlations (Fig. 5.**D**) are lower in dataset 2, indicating that these two parameters are more sensitive to culture conditions. Average growth 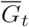 is higher in mutants than in wild-type (Fig. 5.**A**) over the temporal window considered; this might be compensated by lower growth in mutants at later stages or by earlier growth arrest in mutants, because mutant sepals are about 20% smaller in area than wild type sepals [31]. The amplitude of fluctuations Δ*G*_*t*_ is smaller in *spiral2*, but it is not possible to conclude about *katanin*, because the two alleles (*bot1* and *mad5*) show different trends (Fig. 5.**C**). When comparing mutants to wild-type plants, temporal correlations are lower (Fig. 5.**B**), suggesting lower persistence time in mutants. The changes in temporal correlations Γ_*t*_ are lower than in growth rates, so that the changes in non-dimensional persistence time 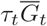 are expected to be dominated by those in growth 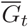, with higher 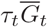 in mutants. This might be ascribed to differences in mechanical responses in these mutants — assuming wild-type plants to have optimal mechanical responses, both over-reaction and under-reaction to mechanical stress would increase the timescale of changes in growth rates [27]. Based on our minimal model of fluctuation stretching (see Eq. 2), smaller non-dimensional persistence time 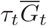 would yield higher extent *α*_*t*_ of spatial correlations. Indeed, the exponent of the Fourier specrum appears higher in mutants (Fig. 5.**D**), although the level of uncertainty makes it difficult to draw a firm conclusion. In the following section, we further test whether fluctuations stretching applies to cell growth in sepals.

### A conserved relation between growth parameters supports fluctuation stretching

We sought relations between growth parameters that would hold across genotypes, data sets, and developmental stages. We first considered the pairwise relations between the growth parameters defined for each sepal: mean growth rate, 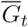, temporal correlation coefficient, Γ_*t*_, normalised amplitude of spatial fluctuations, 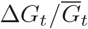, and extent (exponent) of spatial fluctuations, *α*_*t*_. The corresponding scatter plots are shown in Fig. 6.**A-F**. To assess these pairwise relations, we computed Kendall’s correlation coefficient between pairs of parameters. We found rather weak trends overall. The strongest trends were between the exponent, *α*_*t*_, and the temporal correlation coefficient, Γ_*t*_, and between *α*_*t*_ and the average growth 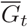. Interestingly, these trends are consistent with fluctuation stretching: larger spatial extent of fluctuations is favored by higher growth rate and by higher temporal persistence, see Eq. 2. We therefore tested more directly the predictions of fluctuation stretching.

**FIG. 6.**
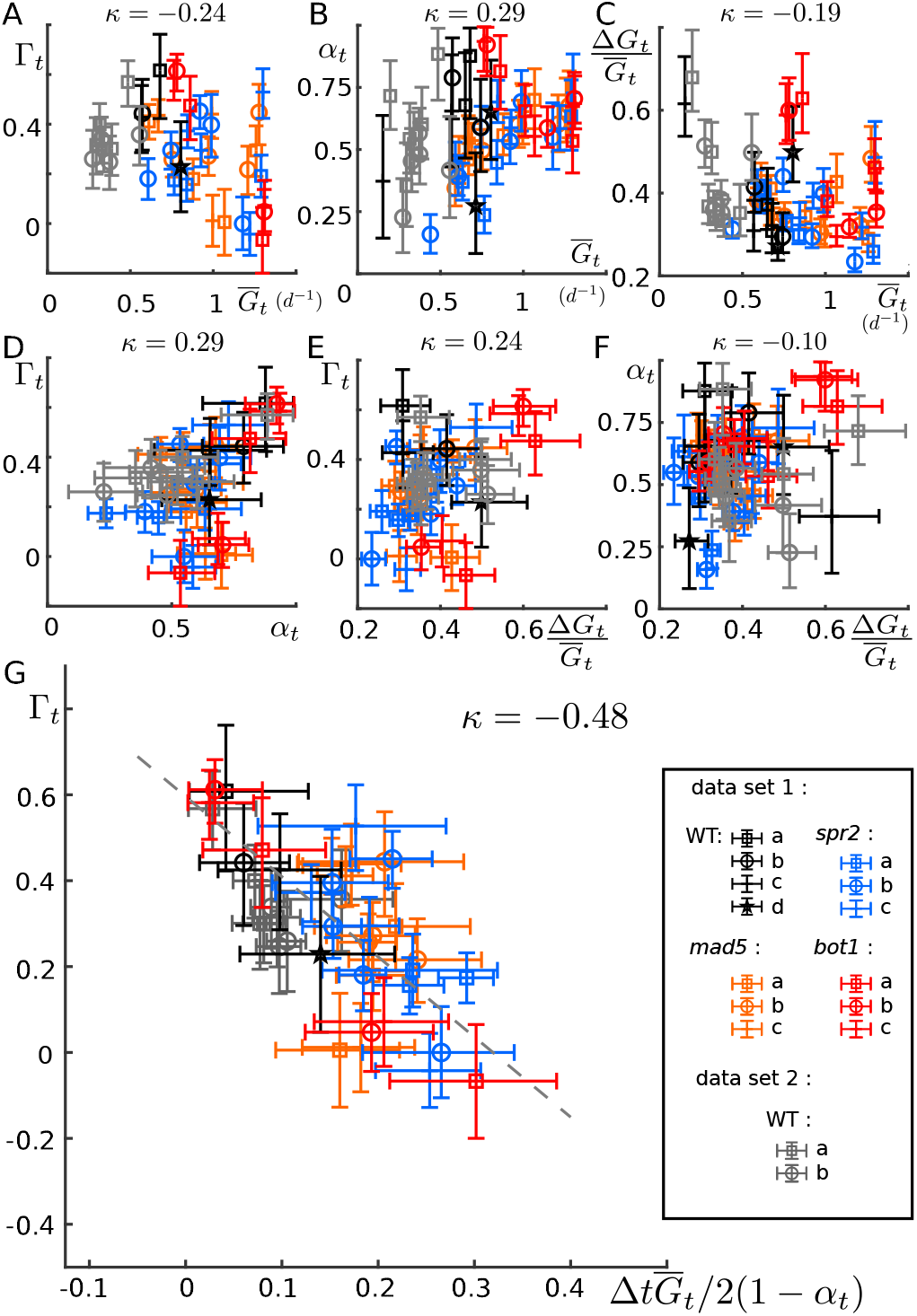
Relations between parameters of growth (fluctuations). **A-F** Pairwise scatter plots of all growth parameters. **A-C** Temporal correlation coefficient Γ_*t*_, exponent of spatial fluctuations *α*_*t*_, and dimensionless amplitude of spatial fluctuations, 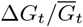, respectively, as function of average growth 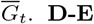 Temporal correlation coefficient, Γ_*t*_, as function of exponent of spatial fluctuations, *α*_*t*_, and dimensionless amplitude of spatial fluctuations, 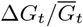, respectively. **F** Exponent of spatial fluctuations, *α*_*t*_, as function of their dimensionless amplitude,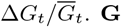. **G** Test of the coupling between temporal and spatial fluctuations, as resulting from fluctuation stretching. Temporal correlation coefficient Γ_*t*_ as a function of the combination 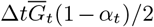 where Δ*t* = 1 *d* is the time step of live imaging. The dashed line corresponds to a linear fit, 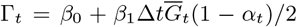, with fit parameters *β*_0_ = 0.596 ± 0.024 and *β*_1_ = −1.87 ± 0.15. The analysis of the fit residuals supports a deterministic relation between the two, see Supplementary note. In all panels, error bars show the 90 % confidence intervals; black, blue, orange, and red symbols correspond to wild-type, *spr2, mad5* and *bot1* sepals from dataset 1, respectively, while gray symbols correspond to wild-type sepals from dataset 2. Kendall’s correlation coefficient, *κ*, is shown above each plot.

Fluctuation stretching does not reduce to a pairwise relation between growth parameters because it relates spatial correlations to time persistence and growth rate. If this phenomenon is at play in sepals, then Eq. 2 and the relation *α* = 1 − *β/*2 (see section Datasets and Methods) imply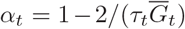, where *τ*_*t*_ is the persistence time. We could measure all parameters of this relation but *τ*_*t*_. Nevertheless the temporal correlation coefficient, Γ_*t*_, should be a decreasing function of Δ*t/τ*_*t*_, Γ_*t*_ = *f* (Δ*t/τ*_*t*_), where *f* is an unknown function and Δ*t* = 1*d* is the time delay between two steps of live imaging, because correlations between states of the sepal at consecutive time steps are higher if the time delay is small compared to the persistence time. By eliminating *τ*_*t*_ from the preceding equations, we found that the time correlation coefficient depends on a combination of the other parameters,

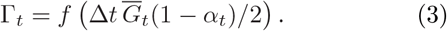

We plotted in Fig. 6**G**. the time correlation coefficient Γ_*t*_ as a function of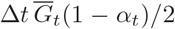. The trend is much clearer than in all other panels of Fig. 6 (Kendall’s coefficient *κ* = − 0.48) and the data seem to collapse along a line. We used statistical inference to perform a linear fit of the data, 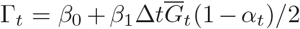, see Supplementary note. We obtained fit parameters *β*_0_ = 0.596 ± 0.024 and *β*_1_ = − 1.87 ± 0.15, with relatively small standard deviations. We then confirmed with a Kolmogorov-Smirnov test that the residuals (the spread of the data around the fit) could be explained by the uncertainty on the estimates of *τ*_*t*_ and Γ_*t*_, see Supplementary note, while the same analysis for the other plots (Fig. 6**A-F**) confirmed that none of these plots was consistent with a linear behavior. Altogether these results support the hypothesis of a deterministic relation between Γ_*t*_ and 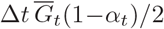 and therefore indicate that fluctuation stretching is at play in growing sepals.

## DISCUSSION

Our analysis provides evidence that growth stretches temporally persistent fluctuations: while no clear pair-wise relation could be made among the different growth parameters, see Fig. 6**A-F**, the clear trend of panel **G** suggest that the persistence time can be deduced from space correlations and tissue growth. This phenomenon explains why higher correlation between cells (higher spatial correlations) may induce more variable organ shape and size [26]. Fluctuation stretching gives a prominent role to the persistence time (correlation time) in controlling spatial correlations in the tissue. Any mechanism that would decrease persistence time would reduce spatial correlations and, as a consequence, variability of organ contours. Accordingly, reducing persistence time would yield robust morphogenesis.

Surprisingly, we found that the temporal correlation coefficient, Γ_*t*_, is generally not much smaller than unity, implying that the persistence time, *τ*_*t*_, is not much smaller than the time scale of growth 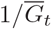. This might be specific to plants. The cell wall sets the local growth rate, and, at the same time, is remodelled at the pace of growth, so that the persistence time of fluctuations of cell wall properties is given by the time scale of growth. It would be worthwhile to extend our study to expanding animal tissues imaged live such as the imaginal disc of the fruit fly [41]. In animal tissues that undergo convergent extension, we would expect fluctuation stretching to operate only in the direction of extension, and so spatial correlations to be highly anisotropic.

As a consequence of fluctuation stretching, the level of time persistence, or more rigorously its product with average growth rate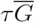, has a strong impact on variability of organ shape and size variability. Indeed, the shape and size of an organ result from the growth of its cells (or of its subcellular elements) integrated over time. If cell growth has a random component, well-defined shape and size may still be obtained through spatiotemporal averaging [26], the cancellation of random effects over large samples (number of cells or time points) — a local excess of growth may be compensated by lower growth later or elsewhere in the tissue. Higher temporal or spatial correlations reduce spatiotemporal averaging since an excess of growth is less likely to be compensated. Accordingly, higher temporal persistence (scaled with growth rate) reduces the robustness of organ shape and size.

We found a higher spatial extent of correlations (higher *α*_*t*_) in mutant genotypes, suggesting higher 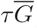. This means that these mutants potentially have more variable shapes or are less robust to perturbations, consistent with the observation that the width of sepals in *bot1* and *spr2* varies more with trichome number in WT plants [31]. We previously predicted that variability of organ contours is minimal for a well-defined level of feedback from mechanical stress to cellulose synthesis [27], leading to the hypothesis that in wild-type sepals the level of mechanical feedback is optimised so as to reduce variability of sepal shape, compared to mutants with lower (*bot1*) or with higher (*spr2*) mechanical feedback. This level of mechanical feedback also corresponds to a minimum of the persistence time of fluctuations (scaled with average growth rate), 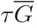, highlighting the importance of this factor in setting the robustness of organ shape and size.

Fluctuation stretching is a kinematic phenomenon: properties of cells or of regions of cells are carried (advected) by tissue growth and deformation; the persistence time of these properties sets how they are carried to larger or smaller spatial scales, in the case of tissue expansion or tissue shrinkage, respectively. This kinematic phenomenon applies to any type of property or field as long as it is carried by tissue growth and deformation, such as protein concentrations in cells. Although fluctuation stretching not only applies to scalar quantities but also to vector fields (e.g. cell polarity) or tensorial fields (e.g. organisation of cytoskeleton), we limited our study to a scalar (areal growth) and did not consider growth anisotropy to avoid the difficulty of taking into account the curved geometry of sepals. Mathematical formalisms such as quasiconformal transforms [42] may nevertheless help to circumvent this difficulty. In the case of complex advective flows, effects associated to co-rotation may arise for non scalar fields. Advection also applies to non-random properties, in line with theoretical models of polarity fields showing that a combination of morphogens, advection, and time persistence can reproduce the shapes of leaves [43], or with models of leaf vasculature that show that areole (region delimited by veins) shape is advected by leaf growth [44].

Altogether, our work sheds light on the role of persistence time, that is the memory of previous states of a given property, in the robustness of morphogenesis. The investigation of spatiotemporal fluctuations may provide a new avenue to characterize organ development.

## AUTHORS CONTRIBUTIONS

Conceptualisation: AF, AB. Data curation: LH, AF. Investigation: AF. Methodology: AF. Writing – original draft: AF, AB. Writing – review and editing: all authors. Supervision: AR, CBL, AB. Funding acquisition: AR, CBL, AB.

## ACKNOWLEDGMENT

We gratefully thank Nathan Hervieux and Olivier Hamant for providing the live imaging data used here. This work was supported by the Human Frontier Science Program grant no. RGP0008/2013 (A.B., A.H.K.R., C.-B.L.), by the US National Institutes of Health Institute of General Medicine grant no. R01 GM134037 (A.H.K.R.), by the US National Science Foundation grant no. MCB-2203275 (A.H.K.R.), by the Université de Lyon through the program “Investissements d’Avenir” grant no. ANR-11-IDEX-0007 (A.B.), and by the French National Research Agency grant no. ANR-21-CE30-0039-01 (A.B.).

## DATASETS AND METHODS

### Model for fluctuation stretching

We introduced a simple model for the dynamics of a quantity Φ(*x, t*) that varies with position vector, *x*, in *D*-dimensional Cartesian space and with time, *t*. We assumed Φ to be advected by tissue growth at rate 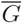, to have a persistence time *τ*, relaxing towards its reference value ⟨Φ⟩, and to be driven by a stochastic source *ξ*(*x, t*), so that

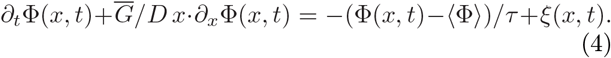

This equation can be solved as shown in the Supplementary note.

### Experimental datasets

In order to reliably analyse fluctuations of growth rate, we chose datasets of sepals imaged with the highest spatial resolution possible among those published. We used live imaging sequences from [32] (dataset 1) and from [31] (dataset 2). Voxel size was 0.12× 0.12× 0.50 *μm*^3^. All plant lines in these sequences were crosses between Ws-4 and Col-0 ecotypes, harbouring respectively the microtubule reporter *p35S::GFP-MBD* and the membrane reporter *pUQ10::Lti6b-2xmCherry* [32]. The two datasets had slightly different culture conditions (type of lighting). Dataset 1 contained wild-type plants, the *spr2-2* allele of *SPIRAL2* that was originally obtained in a Col-0 background, the *bot1-7* allele of *Katanin* that was originally obtained in a Ws-4 background, and the *mad5* allele of *Katanin* that was originally obtained in a Col-0 background (for *mad5*, unpublished sequences were obtained in parallel with those from [32]).

### Segmentation

For sepals not already processed in [31, 32], cells of the abaxial epidermis were segmented and tracked in time using MorphoGraphX [45]. A triangular mesh was obtained for the outer organ surface in which cells were identified and well-delimited.

### Computation of growth rates

We aimed at analysing fluctuations of cell relative areal growth rates tangentially to the sepal and therefore to get rid of the curvature of the outer surface of cells. To do so, we redefined the surface of cells from the linear interpolation of their contours by a flat surface. Areal growth rate was computed from the cell surface area at successive time steps. At time *t*, each cell is labeled by an index *i* and has surface area *S*_*i,t*_. Cell *i* may divide between *t* and *t* + 1; the set *J*_*i,t*_ contains the labels of all daughters of cell *i* at time *t* + 1 (*J*_*i,t*_ is reduced to a single label if cell *i* has not divided). We only consider cells which or whose daugthers remain in the segmented region from *t* to *t* + 1. The areal growth rate of the cell *i* at a time *t* is then defined as

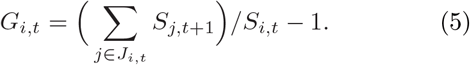

Average (tissular) growth is in turn defined as 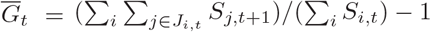.

### Cellular Fourier Transforms

The Fourier harmonics are built from a coarse and discreet version of the Laplace operator. To compute this operator we triangularized cell surfaces using the ‘MESH2D’ matlab algorithm [46, 47]. More details can be found in the Supplementary note. The Cellular Fourier Transform (CFT) 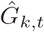 of cell relative areal growth gives the weights by which growth is decomposed over the harmonics *e*_*k*_ of the CFT. In this paper, the definition of the CFT differ from the one in [38] by a prefactor 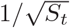 where *S*_*t*_ is the total surface area. This change simplifies the interpretation of Fourier spectra: the coefficients have the same physical dimension as the original signal and the first coefficient is the average of the signal.

### Scaling exponent and amplitude of fluctuations

We quantified spatial correlations in the tissue by fitting the spectral density with a power law. To do so, we assumed a Gaussian distribution for the CFT, centred around 0 with a standard deviation verifying,

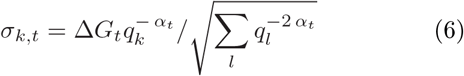

where Δ*G*_*t*_ and the scaling exponent *α*_*t*_ are the fit parameters characterizing respectively the amplitude and the extent of spatial correlation of growth fluctuations. For the fit, we used statistical inference as detailed in the Supplementary note. Doing so, we estimated a probability for the parameters Δ*G*_*t*_ and *α*_*t*_, their expected value, their standard error, and median values. We also estimated the 90% confidence interval, from the fifth to the ninety fifth percentiles.

### Temporal correlations

We estimated temporal correlations of relative areal growth in considering cell growth *G*_*i,t*_ from *t* to *t* + 1 and cells growth 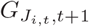 from *t*+1 to *t*+2. 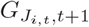 is simply the areal growth rate from *t* to *t* + 1 of the descendants of the cell *i* in the segmentation at *t*:

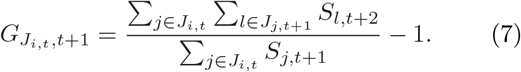

To avoid any bias due to systematic variation of growth at organ scale [32], we used the detrended cell growth *δG*_*i,t*_, which can be defined by subtracting average growth in a local neighborhood from cell growth, see Supplementary note. Temporal correlations were computed as Kendall’s correlation coefficient Γ_*t*_ of *δG*_*i,t*_ and 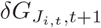. Kendall’s correlation coefficient is rank-based and so is less sensitive to outliers [39]. We used boostrapping to obtain confidence intervals and uncertainties.

We note that Γ_*t*_ tends to be underestimated: A positive error on 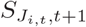 leads to an overestimation of *δG*_*i,t*_ and an underestimation of 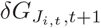, inducing a negative correlation between *δG*_*i,t*_ and 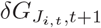. This may explain the few negative values of Γ_*t*_. We found this negative bias to be stronger when we defined growth from the cells outer surface area, leading us to use the interpolation of cell contours instead (see above).

### Comparing genotypes

To describe the impact of mutations or culture conditions on growth parameters, we compared tissues at equivalent developmental stages. We first synchronized all the live imaging sequences from a dataset by building upon the approach developed in [40]. We considered the time curves of organ width for every sepal and finding the time delays ensuring the best superposition between width vs. time curves, leading to a corrected time *T*_*t*_. We checked that this temporal alignment was consistent with stages of guard cell differentiation, indicating that sepal width is a good proxy of developmental stage in the genotypes/conditions that we studied. We defined the mean shift of a quantity Φ_*t*_ as

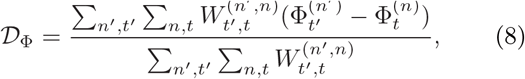

where *n*^′^ and *n* label the pair of sepals compared (e.g. one mutant and the reference wild-type) and *t*^′^ and *t* correspond to the time in the sequence of live-imaging of those two sepals. The sums 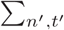 and Σ_*n,t*_ are over all sequences of the mutant and the WT respectively. 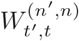 gives the weights at which each pair is considered.

A weight differs from 0 only if the values of synchronized times *T*_*t*_ of the pair are close, see Supplementary note for details. 𝒟_Φ_ quantifies how much, in average, the quantities Φ_*t*_ for the mutants (or for WT in dateset 2) are shifted from the reference WT.

## Supplementary note

### I. MODEL FOR FLUCTUATION STRETCHING

#### A. Model

In line with the explanation of fluctuation stretching proposed in Figs. 1-2, we model the dynamics of a quantity Φ advected in a growing medium. If the medium grows isotropically and uniformly, its strain rate tensor in the *D*-dimensional space is 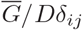 where 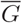 is the line, surface or volume growth for *D* = 1, 2, or 3 respectively and *δ*_*ij*_ is the Kronecker delta tensor. We assume the dynamics of the quantity Φ to be ruled by intrinsic cellular processes among which some are stochastic. For simplicity, we restrict our model to lowest order and consider a linear partial differential equation. Denoting time by *t* and the Cartesian space coordinate vector by *x*, we assume the evolution of Φ(*x, t*) to be given by

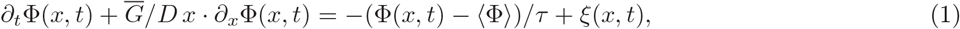

where the material point at *x* = 0 serves as the origin of the spatial coordinate system. *∂*_*t*_ and *∂*_*x*_ respectively stand for the partial derivative with respect to time and for the gradient. The left hand side of (1) corresponds to the material time derivative. The first term in the right hand side ensures the relaxation of Φ toward its reference value ⟨ Φ ⟩ with a time scale *τ*, while the second term accounts for stochasticity through the noise *ξ*.

#### B. Linear response

We denote the deviation of Φ from its reference value by ΔΦ(*x, t*) = Φ(*x, t*) − ⟨ Φ ⟩. The persistence time *τ* sets the memory of the system as can be seen in the explicit solution of (1),

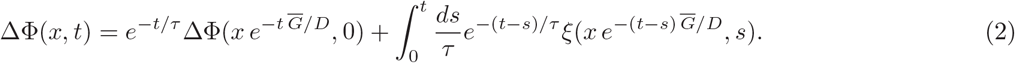

In this equation, *τ* sets the time over which initial conditions persist and the delay over which the noise impacts the value of Φ.

#### C. Spatial correlation function

To describe the statistical properties of Φ, we assume the noise to be Gaussian, with ⟨ *ξ*(*x, t*) ⟩ = 0 and ⟨ *ξ*(*x, t*)*ξ*(*x* + *l, t* + *s*) ⟩ = *Kδ*(*s*)*g*( | *l* |). ⟨ · ⟩ stands for an ensemble average, *K* is the noise strength, and *δ*(·) is the Dirac distribution. The function *g*(*l*) = ⟨ *ξ*(*x, t*)*ξ*(*x* + *l, t*) ⟩ */* ⟨ | *ξ*(*x, t*) | ^2^ ⟩ describes the spatial correlations of *ξ*, assumed to be regular and to vanish at infinity. As a consequence of the long-ranged correlations that we predict, small scales cannot be neglected and a Dirac distribution cannot be substituted to *g* without causing problems of convergence, unless a cutoff is introduced by hand. The correlations of Φ can be computed using (2) with *t* = −∞ as initial time. The space correlation function *C*(*l*) = ⟨ ΔΦ(*x, t*)ΔΦ(*x* + *l, t*) ⟩*/* ⟨ |ΔΦ(*x, t*)|^2^ ⟩ can then be written as,

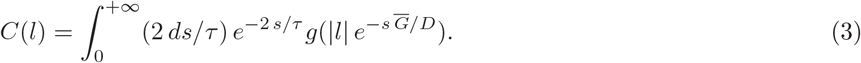

The space correlation function *C*(*l*) is obtained by stretching the variation lengthscales of *g* by a factor 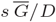 and summing the stretched functions with weights *e*^−2*s/τ*^. Changing the integration variable, we rewrite (3) as,

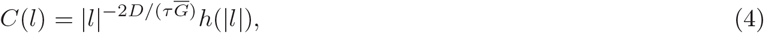

where the increasing function 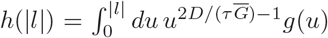 is expected to reach an asymptotic value as | *l* | is large compared to the correlation length of *ξ*. (4) makes therefore explicit the long-ranged property of *C*, characterized by the scaling exponent 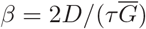.

#### D. Fourier spectrum

The Fourier transform 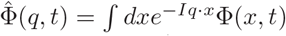 can be used to estimate the space correlation function *C*(*l*). More exactly, the mean squared spectrum 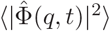 is proportional to the Fourier transform 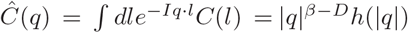 with 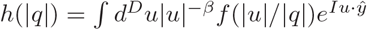 and *ŷ* a unit vector. It exhibits a singularity for | *q* | → 0 where it scales like | *q* |^−2*α*^, with

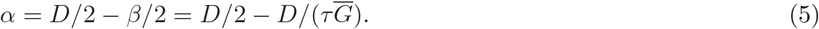

If the correlation length of the noise source is small with respect to system size, the root mean squared spectrum can be approximated by a power law whose amplitude relate to the standard deviation through Parseval’s theorem and whose exponent *α* is given by the persistence time *τ* and growth rate 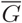 according to (5).

### II. CELLULAR FOURIER TRANSFORM

Here we present the computation of cell surface area, we define the discrete Laplace operator, we explain how we built the Fourier harmonics based on this Laplace operator, and we define the Cellular Fourier Transform (CFT). The theoretical basis of the CFT may be found in [1].

#### A. Cell area and discrete Laplace operator

We compute cell area from the linear interpolation of cell contour. More precisely, we project the contour on a plane that is perpendicular to the surface vector. The contour being polygonal, the surface factor can be written 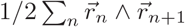 where the sum is over the contour vertexes, 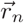 is their position, *n* indexes the position around the contour and is the exterior product. We then triangulate the surface enclosed in the projected contour using the MESH2D Matlab package [2, 3]. To obtain a 3D mesh and determine the position of the mesh along the surface vector, we performed a linear interpolation of the cell contours. The area *S*_*i,t*_ for cell *i* at time *t* is then computed as the sum of areas of triangles in the triangulation, 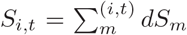, where *m* spans triangles of cell *i* at time *t* and *dS*_*m*_ is the area of triangle #*m*. The tissue is made of *N* cells that are followed from *t* to *t* + 1.

The discrete Laplace operator is a square matrix of size *N* × *N* and its components are given by

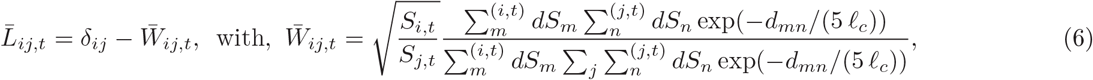

where indices *i* = 0, 1, ..*N* − 1 and *j* = 0, 1, ..*N* − 1 span the *N* cells of the tissue. *d*_*mn*_ is the distance between triangle *m* from cell *i* and triangle *n* from cell *j*, both considered at time step *t*. The unit of length is mean cell size 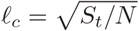, where *S*_*t*_ is the surface of the tissue at time *t* and *N* is the number of cells. Here we took the width 5*l*_*c*_ for the coarse Laplace operator.

#### B. Fourier harmonics

We define Fourier harmonics as the right singular vectors of the discrete Laplace operator 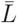 defined in Eq. 6. We showed in [1] that 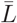 is a good representation of the coarse Laplace operator *ℒ* [*f* ](*x*) = *dy* exp (|*x* − *y*|*/*(5 *𝓁*_*c*_)) (*f* (*x*) − *f* (*y*)), applying to real functions *f* of the position vector. The singular vectors of 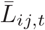 are, for example, expected to have the same oscillatory nature as the eigenfunctions of *ℒ* and their associated wave number *q*_*k*_ to relate to their singular values through the same relation *q*_*k*_ = 1*/*(5*l*_*c*_)*Q*(*λ*_*k*_), with 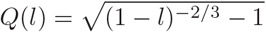 associated to the kernel of the coarse Laplace operator [1]. The singular value decomposition of the Laplace operator 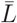, which yields left singular vectors *V*, right singular vectors *U*, and the singular values 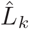, is:

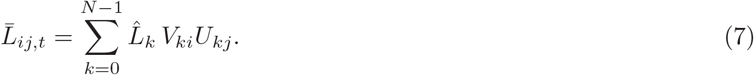

The value taken by the *k*^th^-harmonic in cell *i* at time step *t* is 1*/S*_*i,t*_*U*_*ki*_, and its wave number is given by 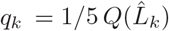. The harmonics are indexed so that their index grows with the wave number.

#### C. Calculation of the CFT of cell growth

The areal growth rate of cell *i* at time step *t* is defined as 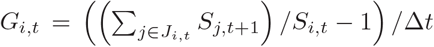 where *J*_*i,t*_ is either the new label of cell *i* at time *t* + 1 or the set of labels of the daughters of cell *i* if it has divided, while the time step is always Δ*t* = 1*d*. The *k*^th^ CFT coefficient is then 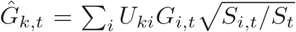 where *S*_*t*_ is the total area *S*_*t*_ = ∑_*i*_ *S*_*i,t*_. Here we use a convention that differs from [1] by a multiplicative factor 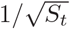 in the definition of the CFT. This makes the interpretation of CFTs simpler: they have the same dimensions (units) as the original signal (here growth) and the first coefficient is equal to the average signal.

### III. SPATIAL CORRELATIONS

We estimated spatial correlations of growth from the Fourier spectra *i*.*e*. from the distribution of Fourier transforms 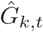 and associated wavenumbers *q*_*k*_. For this we used Bayesian inference.

#### A. Inference methods applied to Fourier spectra

To quantify spatial correlations, we assumed the CFT coefficients, 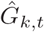 for *k* ≥ 2, to be independent random Gaussian variables whose mean squared deviation follows a power law with respect to the wave number *q*_*k*_,

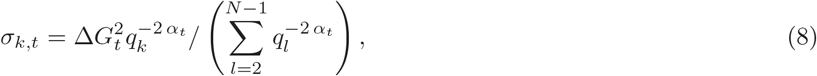

with the parameters Δ*G*_*t*_ and *α*_*t*_ quantifying the amplitude of growth fluctuations and their space correlations, respectively. We made the choice not to consider the first two CFT coefficients to avoid potential bias related to large scale growth patterns, which should not be considered as fluctuations. For the derivation of the equations, it is more convenient to rewrite (8) as 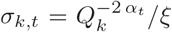, where 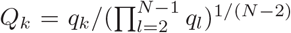 and 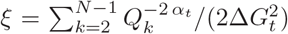. We write the probability distribution fucntion of 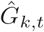 as

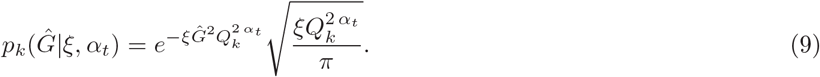

We use Bayesian inference to estimate *ξ* and *α*_*t*_, assuming a flat prior distribution for *ξ* ∈ [0, + ∞ [ and *α*_*t*_ ∈ [0, 1], which are the relevant range of parameters for (9). The posterior distribution for *ξ* and *α*_*t*_ takes the form

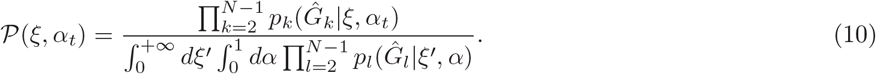

We then substitute the probabilities *p*_*k*_ by their explicit form, noting that, by construction, 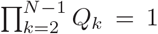, and, computing the first integral in the denominator, we get

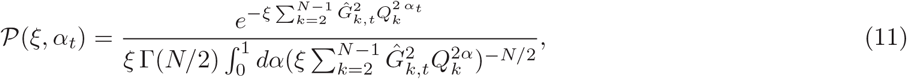

where Γ is Euler’s gamma function.

#### B. Estimating amplitude of fluctuations and exponent of spatial correlations

To estimate Δ*G*_*t*_, *α*_*t*_ and their uncertainty, we consider the joint cumulative distribution function *ℱ* (Δ*G, α*), of having Δ*G*_*t*_ and *α*_*t*_ smaller than the values Δ*G* and *α*, respectively. This function can be written in terms of *𝒫* (*ξ, α*) as

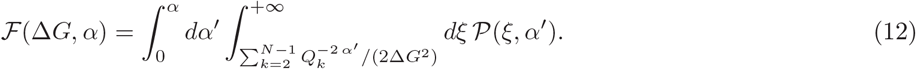

By using the expression *𝒫* (*ξ, α*) in (11) and computing the second integral, we then get

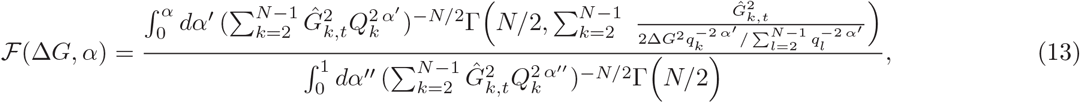

where 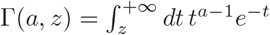 is the incomplete gamma function.

We used the median as a representative value of the different quantities we considered. We estimated Δ*G*_*t*_ from the median *ℱ* (Δ*G*_*t*_, 2) = .5 and the 90% confidence interval [Δ*G*_1,*t*_, Δ*G*_2,*t*_] from the 5^th^, *ℱ* (Δ*G*_1,*t*_, 2) = .05, and the 95^th^ percentile, *ℱ* (Δ*G*_2,*t*_, 2) = .95. Similarly, we estimate *α*_*t*_ from the median *ℱ* (+ ∞, *α*_*t*_) = .5 and the 90% confidence interval [*α*_1,*t*_, *α*_2,*t*_] from the 5^th^, *ℱ* (+ ∞, *α*_1,*t*_) = .05, and the 95^th^ percentile, (+ ∞, *α*_2,*t*_) = .95.

When we approximated their distributions by Gaussians (for fits or to estimate shifts from WT to mutants tissues), we used the the expected value and the standard deviations of *α*_*t*_ and Δ*G*_*t*_. We estimated the expected value of *α*_*t*_,

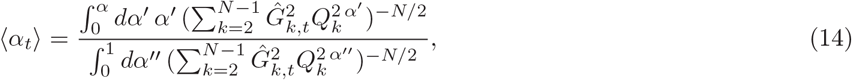

its standard deviation 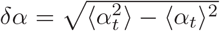 with,

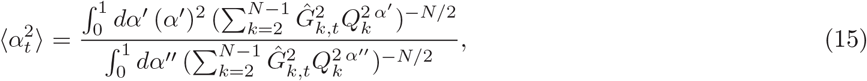

the expected value of Δ*G*_*t*_,

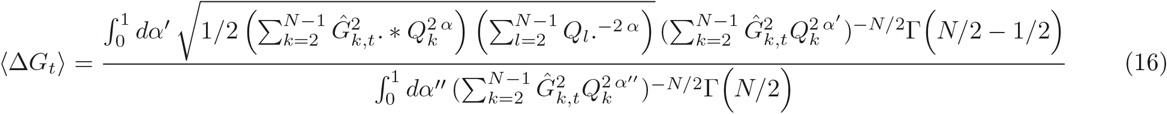

and the standard deviation 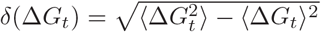

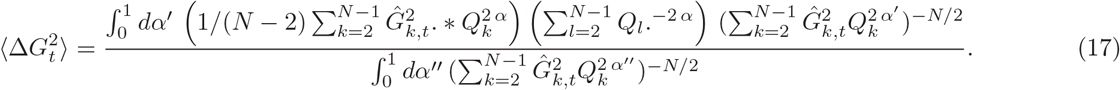

### IV. TEMPORAL CORRELATIONS

To quantify temporal correlations, we detrended growth from large-scale spatial patterns and we calculated Kendall’s correlation coefficient of detrended growth.

#### A. Detrending

Before estimating time correlations, we corrected cellular growth using a local average of growth, aiming to detrend our estimate from large-scale deterministic spatial variations. We thus avoid potential bias induced by large scale growth variations that should not be considered as fluctuations. We use growth rate *G*_*i,t*_ of cell *i* between *t* and *t* + 1, as defined in Sec. II A. Computing local excess of growth is equivalent to apply a smooth Laplace operator to growth [1]. For convenience, we use the Laplace operator defined in (6), and we define 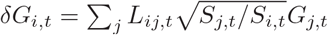, where *j* spans cells that can be tracked from *t* to *t*+2. Detreneded growth at time *t* needs to be compared to detrended growth at time *t* + 1, 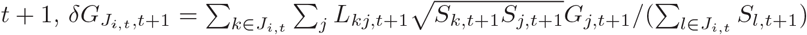.

#### B. Kendall’s correlation coefficient

Time correlations are quantified by Kendall’s correlation coefficient Γ_*t*_ between *δG*_*i,t*_ and 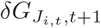. We used a bootstrap approach with 10^4^ resamplings to quantify the statistical properties of Γ_*t*_. We estimated Γ_*t*_ from the median of the boostrap distribution and the bounds of the confidence interval are its 5^th^ and its 95^th^ percentile. Finally, we also considered ⟨Γ_*t*_ ⟩ and *δ*Γ_*t*_ the expected value and the standard error of the distribution.

### V. ANALYSIS OF TEMPORAL VARIATIONS IN GROWTH PAREMETERS

We analyzed two datasets, the first containing wild-type and mutant plants while the second group contained wild-type plants grown in different conditions. We first synchronised the time series of the two datasets. We then compared mutants to wild-type sepals from plants cultured in the same conditions, or wild type sepals from plant cultured in different conditions.

#### A. Registration

To synchronize (register) the different time series (labeled with an upper index (*n*)), we looked for the temporal shifts Δ*t*^(*n*)^ that maximise the overlap of curves of width vs. time 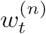. The perfect overlap being, in general, not possible, we define a distance between pairs of curves, and we choose the delays which minimise the quadratic sum over all possible pairs 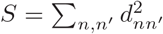, of these distances. For two time series 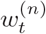 and 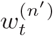, the distance from *n* to *n*^′^ is defined as 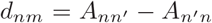, where 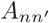 is the area of the region in the Cartesian plane that is delimited to the left by the linear interpolation of 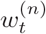 versus *t* and to the right by the linear interpolation of 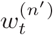 versus *t*. This distance depends linearly on the the time-shifts, 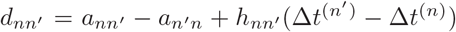 and 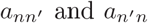 are the areas 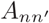 and 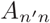 before synchronization. The minimization problem is then simply quadratic and the shifts are given by the solution of

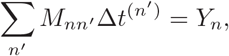

with 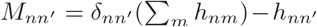 and 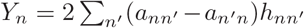. The matrix *M* is not invertible due to invariance by translations in time, but this system can be solved by adding the condition that the smallest temporal shift (the smallest Δ*t*^(*n*)^) has a value of 0. We denote by 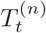 the new temporal coordinate for live-imaging series *n* following registration. We checked that this temporal alignment was consistent with stages of guard cell differentiation, indicating that sepal width is a good proxy of developmental stage in the genotypes/conditions that we studied.

#### B. Differences between mutant and wild-type growth parameters

To compare a quantity Φ_*t*_ (which could be Γ_*t*_, Δ*G*_*t*_, *α*_*t*_ or 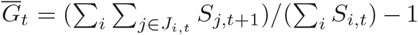) between mutant sepals or wild-type from dataset 2 and wild-type sepals from dataset 1, we defined the mean difference *𝒟*_Φ_ as,

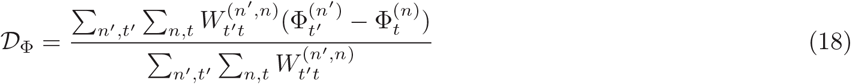

where the upper indices (*n*^′^) and (*n*) label the mutant and wild-type live-imaging sequences, respectively. The sums 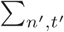 and 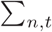 are over all the time points of the mutant and the wild-type, respectively. *𝒟*_Φ_ quantifies how much, on average, the quantities Φ*t* for the mutants differ from the WT. The weights 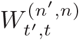 are defined as

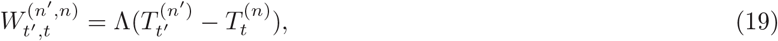

where Λ(*x*) = max(1 − |*x*|, 0) is the triangle function. This definition ensures that only differences between sepals of comparable stages are considered in the distance *𝒟*_Φ_.

Approximating the distribution of Φ_*t*_ to Gaussian, *𝒟*_Φ_ has a Gaussian distribution and its expected value is

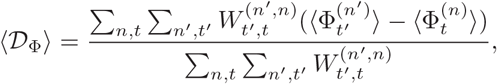

where 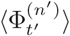 and 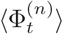 are the expected values of Φ for the mutants and the wild-type tissues. The standard deviation is

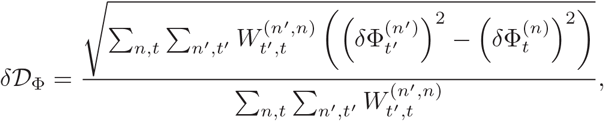

where 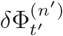 and 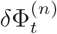 are the standard error of Φ for the mutants and the WT tissues.

### VI. LINEAR FIT AND RESIDUALS

We used statistical inference to determine which linear relation is the most likely to fit our data. We did this to test if the master curve of Γ_*t*_ as function Δ*t/τ*_*t*_ can well be fitted by a linear relation. We also estimated the uncertainty of the fit itself and tested whether the distribution of data around the fit can be explained by the data uncertainty, in coherence with the hypothesis of a linear and deterministic relation between the two.

#### A. Linear fit

We performed this analysis to fit the master curve Γ_*t*_ as function of Δ*t/τ*_*t*_, but since we applied the same analysis to other scatter plots, we considered here the relation between generic variables, *x* and *y*. To each measurement performed (indexed *i*) is associated a probability *p*_*i*_(*x*_*i*_, *y*_*i*_) of finding a certain quantity *x*_*i*_ associated to the quantity *y*_*i*_. Approximating *p*_*i*_ to a Gaussian distribution, and assuming no specific correlations for the error on *x*_*i*_ and *y*_*i*_, we can write

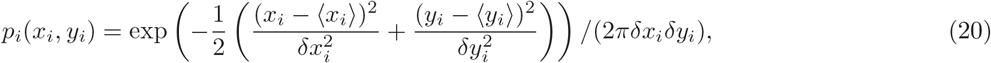

where ⟨ *x*_*i*_ ⟩ and ⟨ *y*_*i*_ ⟩ are the expected values of *x*_*i*_ and *y*_*i*_ and *δx*_*i*_, and *δy*_*i*_ are their standard errors. The probability of finding the *x*-coordinate in *x*_*i*_ and of being on the line *y* = *β*_0_ + *β*_1_*x* is then, *p*_*i*_(*x*_*i*_, *β*_0_ + *β*_1_*x*_*i*_) which can be written as

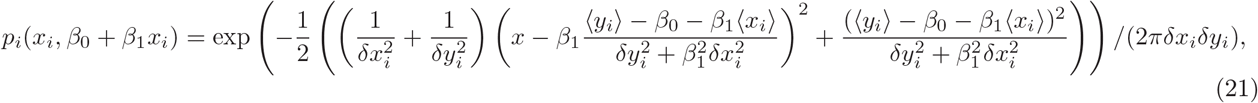

where we rearranged the argument of the exponential to write the dependence with *x* as a square. Integrating over *x*_*i*_, we obtain the probability that the data measured in *i* falls on the line *y* = *β*_0_ + *β*_1_*x* as

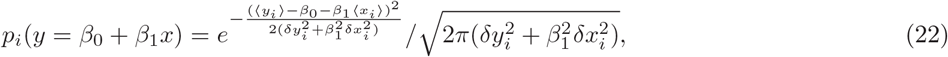

The probability of having the *n*, assumed independent, measurements falling on *y* = *β*_0_ + *β*_1_*x* is then 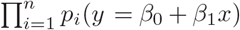, and using flat prior for *β*_0_ and a Cauchy distribution as a prior for *β*_1_, which is equivalent to assume a flat prior for the orientation of the line *y* = *β*_0_ + *β*_1_*x*, we get

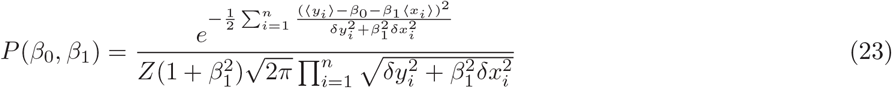

where the constant *Z* given below is defined so that 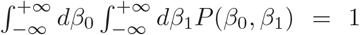. Introducing 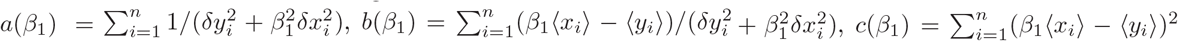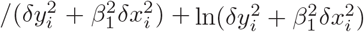, we can write

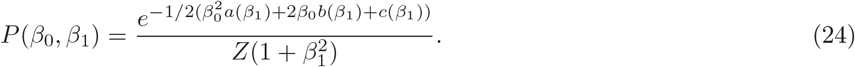

Then, 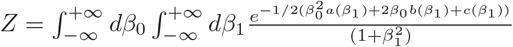 can be rewritten, computing the first integral, as

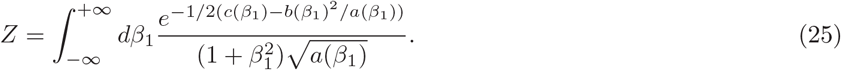

The expected value for *β*_1_ is thus

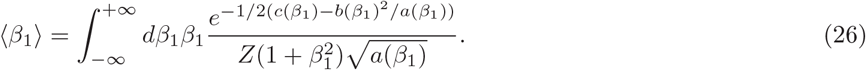

and the standard deviation is 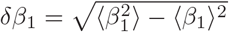, where

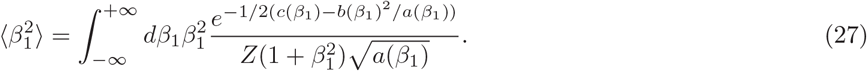

The expected value for *β*_0_ is

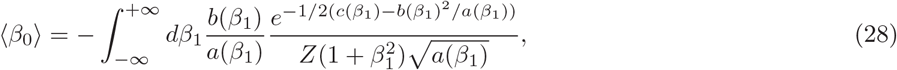

and the standard deviation 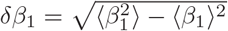, where

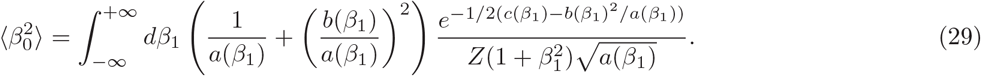

We computed these integrals numerically to estimate the fitting parameters and their standard deviations.

#### B. residuals

We would like to test whether the expected values ⟨ *β*_0_ ⟩ and ⟨ *β*_1_ ⟩ enable to adequately fit the set of data. We gave in Eq. 22 the probability of having a linear relation *y* = *β*_0_ + *β*_1_*x* in measurement *i*. For *β*_0_ = ⟨ *β*_0_ ⟩ and *β*_1_ = ⟨ *β*_1_ ⟩, this probability is

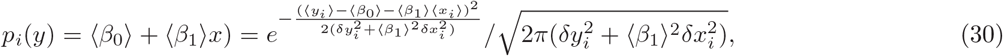

We see that this probability follows a standard normal distribution with respect to the parameter 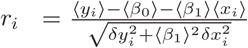. If our assumptions are consistent, and notably the assumption that a linear relation exists between *y*_*i*_ and *x*_*i*_ is correct, then the distribution of *r*_*i*_ over all the measurements should be close to a standard normal distribution. To assess this, we performed a Kolmogorow-Smirnov test at the 5% significance level. We concluded that, in the case of the master curve, the distribution of data around the fit can be explained by the uncertainty on the estimates, and that the data are compatible with the hypothesis of a linear and deterministic relation between Γ_*t*_ and Δ*t/τ*_*t*_, while we could not draw the same conclusions for any of the other pairwise trends. The p-values of the Kolmogorow-Smirnov test for the residuals of the linear fits of all the plots of Fig. 6. of the main are given in the table below.

**TABLE 1.** p-value for the Kolmogorow-Smirnov test of the residuals of the linear fits of all the plots in Fig. 6.

| Plot | p-value |
| --- | --- |
| $\Gamma_t$ vs $\overline{G}_t$ | $2.2 \cdot 10^{-4}$ |
| $\alpha_t$ vs $\overline{G}_t$ | $4.3 \cdot 10^{-2}$ |
| $\Delta G_t/\overline{G}_t$ vs $\overline{G}_t$ | $3.3 \cdot 10^{-7}$ |
| $\Gamma_t$ vs $\alpha_t$ | $2.0 \cdot 10^{-2}$ |
| $\Gamma_t$ vs $\Delta G_t/\overline{G}_t$ | $8.7 \cdot 10^{-3}$ |
| $\alpha_t$ vs $\Delta G_t/\overline{G}_t$ | $1.6 \cdot 10^{-5}$ |
| $\Gamma_t$ vs $\Delta t G_t/2(1 - \alpha_t)$ | $1.1 \cdot 10^{-1}$ |

